# No evidence for paternal age effects on sons or daughters, when accounting for paternal sperm storage

**DOI:** 10.1101/2024.09.19.613916

**Authors:** Krish Sanghvi, Samuel Gascoigne, Biliana Todorova, Regina Vega-Trejo, Tommaso Pizzari, Irem Sepil

## Abstract

1. The age at which a father reproduces is predicted to affect not only his own fertility but also the fitness of his offspring. Specifically, offspring born to old fathers are assumed to be of a lower quality than those conceived by young fathers. However, when fathers have low mating rates, paternal age might be confounded with the duration for which mature sperm are stored in fathers prior to ejaculation.
2. Studies that disentangle the confounding effects of paternal sperm storage duration from those of paternal age, on offspring, are lacking. We use *Drosophila melanogaster* to test the separate and interactive effects of paternal age and sperm storage duration (sexual rest) on the survival and lifetime reproduction of offspring.
3. As expected, old fathers produce fewer offspring than young fathers. But surprisingly, paternal age does not influence the survival or lifetime reproductive success of either sons or daughters. Instead, sons conceived by fathers with long durations of sexual rest have lower reproductive success than sons conceived by fathers with short durations of sexual rest.
4. We further discover that daughters of low reproductive quality selectively disappear with age, but sons do not, highlighting that demographic processes need to be accounted for when studying paternal age effects.
5. Overall, our study suggests that paternal age effects might not be as pervasive as previously assumed. We emphasize that a more nuanced understanding of the mechanisms causing paternal age effects is required, so that studies do not misattribute effects of paternal sperm storage to age.

## Introduction

The influence of parental environments is not limited to the reproductive success of parents themselves. Parental environments and phenotypes can influence offspring phenotypes (Badyaev and Uller, 2009; Liu and Chen, 2018) via genetic (e.g. mutation accumulation in gametes) or epigenetic mechanisms (Bauch et al, 2019; Chen et al, 2016; Heidinger et al, 2016; Perez and Lehner, 2019; Rando, 2016; Rodgers et al, 2015; Sharma, 2019; Yoshizaki et al, 2021), as well as differential resource allocation by parents (Uller, 2008). One parental effect that has received considerable attention is the ‘paternal age effect’. Paternal age effects are caused when the age at which a father conceives offspring affects the offspring’s phenotype (Monaghan and Metcalfe, 2019). These effects have broad implications for healthspan (Chan and Robaire, 2022), life-history evolution, and population dynamics (Evans, et al, 2019).

Paternal age has been hypothesized to be more potent in influencing offspring phenotypes than maternal age (de Manuel et al, 2022; Gao et al, 2019). This rationale stems from sperm producing more reactive oxygen species, but have poorer DNA repair machinery than eggs, and male germlines accumulating more mutations with advancing age than female germlines (Crow, 2000; Ellegren, 2007; Girard et al, 2016; Reinhardt and Turnell, 2000; Venn et al, 2014). Evidence for deleterious paternal age effects includes old fathers producing offspring with poorer development (e.g. Janecka et al, 2017; Preston et al, 2015), lower juvenile survival (e.g. Fay et al, 2016), reduced adult lifespans (e.g. Crow, 2003; Noguera et al, 2018; Priest et al, 2002; Sharma et al, 2015; Wylde et al, 2019; Xie et al, 2018), and lower reproductive output (e.g. Arslan, 2017; Schroeder et al, 2015; Vuarin et al, 2021), compared to offspring conceived by young fathers.

In some cases, the effects of paternal age can be confounded by the duration for which sperm are stored post-meiosis and prior to ejaculation in males (Pizzari et al, 2008; Reinhardt, 2007; Siva-Jothy, 2000). These confounding effects typically arise in studies where fathers are kept virgins for long durations (reviewed in Sanghvi et al, 2024), have low rates of sperm loss or resorption, and have life-long spermatogenesis (e.g. most vertebrates and some insects: Bjork et al, 2007; Demarco et al, 2014; Reinhardt et al, 2011; Santos et al, 2023; Sepil et al, 2020). In such cases, old, virgin males not only have a more senescent germline, but also store sperm for longer durations compared to young virgin males, due to longer periods of sexual rest (Pizzari et al, 2008). Differences in durations of sperm storage between old and young fathers can also arise when fathers of different ages differ in their mating rates (Aich et al, 2022). In natural settings for instance, old males might be mating less often, thus storing sperm for longer durations on average, than young males.

Sperm storage post-meiosis and before ejaculation can affect sperm quality, thus the fertility of a male, as well as his offspring’s fitness, independent of the male’s age (Pizzari et al, 2008, Reinhardt, 2007). Prolonged storage of sperm in males or females can deteriorate sperm quality (Brindle et al, 2023; Cattelan and Gasparini, 2021; Comar et al, 2017; Gasparini et al, 2014, 2019; Hettyey et al, 2012; Levitas et al, 2005; Radhakrishnan and Fedorka, 2011), increase the number of mutations in sperm (Agarwal et al, 2016; Rinehart, 1969), and reduce male fertilisation success (Gasparini et al, 2018; Reinhardt and Siva-Jothy, 2005). Sperm storage independently can also negatively affect the development (Dharmarajan, 1950; Lodge et al, 1971; Pineaux et al, 2019; White et al, 2008), quality (Tarin, 2000; Wagner et al, 2004; White et al, 2008), and fertility (Gasparini et al, 2017) of resultant offspring. This deterioration of stored sperm mainly occurs because sperm accumulate DNA and oxidative damage over prolonged periods of sexual rest (Barbagallo et al, 2022; Sorensen et al, 2023; Wetzker et al, 2024).

When the effects of male age and sexual rest cannot be disentangled, it remains unclear how these processes independently influence male reproductive output. Studies that disentangle these effects report contrasting results to each other (Gasparini et al, 2019; Vega- Trejo et al, 2019). Furthermore, no study has yet tested the separate effects of paternal age and sexual rest on the reproductive output of the conceived offspring. Previous studies that measure offspring phenotypes focus mainly on offspring survival (Meunier et al, 2022), which might not be informative of offspring fitness when offspring survival and reproduction co-vary negatively. Testing the separate effects of paternal age and sperm storage duration is necessary to ensure that studies are not incorrectly attributing the effects of paternal sexual rest to paternal age. Paternal age and sperm storage duration might also interact (Pizzari et al, 2008), for example, if old fathers are worse at repairing damage in stored sperm, than young fathers (Gorbunova et al, 2007; Selvaratnam et al, 2015; Weirich-Schwaiger et al, 1994).

Studies that manipulate both, paternal sperm storage duration/sexual rest and paternal age, and simultaneously measuring offspring and paternal fitness components, are lacking, leaving such hypotheses untested.

Here, we experimentally test the independent and interactive effects of paternal age at conception and paternal duration of sexual rest (henceforth, sperm storage duration), on paternal fertility, and the age-dependent survival and lifetime reproduction of their sons and daughters. We use the fruit fly, *Drosophila melanogaster,* a model organism for investigating parental age effects (Aguilar et al, 2023; Hercus and Hoffmann, 2000; Mossman et al, 2019; Nystrand and Dowling, 2014; Price and Hansen, 1998; Sanghvi, Pizzari et al, 2024; Sepil et al, 2020; Tan et al, 2013), owing to their short generation time and absence of parental care. Fruit flies show life-long spermatogenesis (e.g. Bjork et al, 2007; Sepil et al, 2020) and low rates of sperm loss (Demarco et al, 2014; Santos et al, 2023), leading to age-dependent accumulation of mature sperm in unmated males (Pisano et al, 1993; Sepil et al. 2020). In fruit flies, sperm storage reduces the viability of sperm by 50% over 8 days (Radhakrishnan and Fedorka, 2011), increases oxidative stress by 10% (Wetzker et al, 2024), leads to sperm ejection by females (Snook and Hosken, 2004), and lowers offspring viability (Tan et al, 2013).

We test predictions of five hypotheses (H1-H5). First, paternal sperm storage could interact with paternal age, to influence the reproductive output of fathers, and the phenotypes of their offspring. If previous studies that do not separate these effects, are misattributing the effects of paternal sperm storage to paternal age, we predict that separating these effects will reveal no significant evidence for reproductive senescence in fathers, or for paternal age effects on offspring. Second, old males might be worse at repairing cellular damage than young males (Gorbunova et al, 2007; Weirich-Schwaiger et al, 1994; Witt et al, 2023), causing deleterious effects of sperm storage to be exacerbated in old than young fathers (Zubkova and Robaire, 2006). We thus predict offspring of old fathers with long durations of sexual rest will have lower fitness than offspring of young fathers or of fathers with short sperm storage durations (H2). Third, offspring of old fathers might inherit a higher mutation load than offspring of young fathers (Chen et al, 2023; de Manuel et al, 2022; Girard et al, 2016; Jonsson et al, 2017; Kong et al, 2012; Wang et al, 2020). If paternally inherited mutations are exacerbated as offspring grow older (Bregdahl et al, 2020, 2023; Monaghan et al, 2020; Moorad and Promislow, 2008; Shindyapina et al, 2020), we predict that the effects of old paternal age might be more deleterious when offspring are old than young (H3).

Fourth, paternal age effects might be sex-specific (e.g. Aich et al, 2022; Angell et al, 2022; Gasparini et al, 2017; Krishna et al, 2012; Sparks et al, 2022). For instance, advancing paternal age might deteriorate the Y chromosome (Byrne et al, 2003; Carothers et al, 1978) or imprinted genes (Denomme et al, 2020; Paczkowski et al, 2015). Similarly, telomeres or epigenetic markers (which are affected by paternal age) might be sex-specifically inherited (Bouwhuis et al, 2015; Olsson et al, 2011; Schroeder et al, 2015). In line with these hypotheses, we predict that paternal age and paternal sperm storage duration will have a larger impact on sons compared to daughters fitness (H4). Lastly, fathers might trade-off investment in sperm production versus repair/maintenance of sperm quality (e.g. Koppik et al, 2023). This trade-off between investment in sperm production versus sperm repair might manifest as trade-offs between offspring quantity and quality, therefore modulate paternal age effects (Fischer et al, 2011; Ratikainen et al, 2018; Johnson et al, 2018). In line with these hypotheses, we predict that fathers who produce more offspring would produce poorer quality offspring (H5).

## Replication statement

| Scale of inference | Scale at which factor of interest is applied | Number of replicates at the appropriate scale |
| --- | --- | --- |
| Species | Individuals within blocks | 60 fathers per treatment across 4 blocks. 3 sons and 3 daughters per father measured |

### Methods

#### Stock and experimental individuals

We used *Dahomey* wild-type *Drosophila melanogaster* flies maintained at a 12:12hr light cycle, at a constant temperature of 25°C, and fed with Lewis medium (Lewis, 1960) supplemented with *ad libitum* live yeast (following Sepil et al, 2020). The stock from which these flies were taken have been reared in the lab since the 1970s. Under these conditions, flies have an egg-to-adult development of ∼10 days, and virgin adult males have median and maximum lifespans of ∼45 and 90 days, respectively (Sepil et al, 2020; also see Fig S1). Our experiment consisted of two parts: first, fathers were generated and assigned across four treatments (old or young paternal age, with long or short sperm storage duration) in a fully balanced design (henceforth “F0 assays”). Then, sons and daughters (F1 individuals) from fathers in each paternal treatment were collected, and their survival and lifetime reproduction measured (henceforth “F1 assays). Our experiment (Figure 1) was conducted using a total of 60 fathers per treatment (our independent sample size), spread across four replicates (see Table S1 for sample sizes).

**Figure 1:**
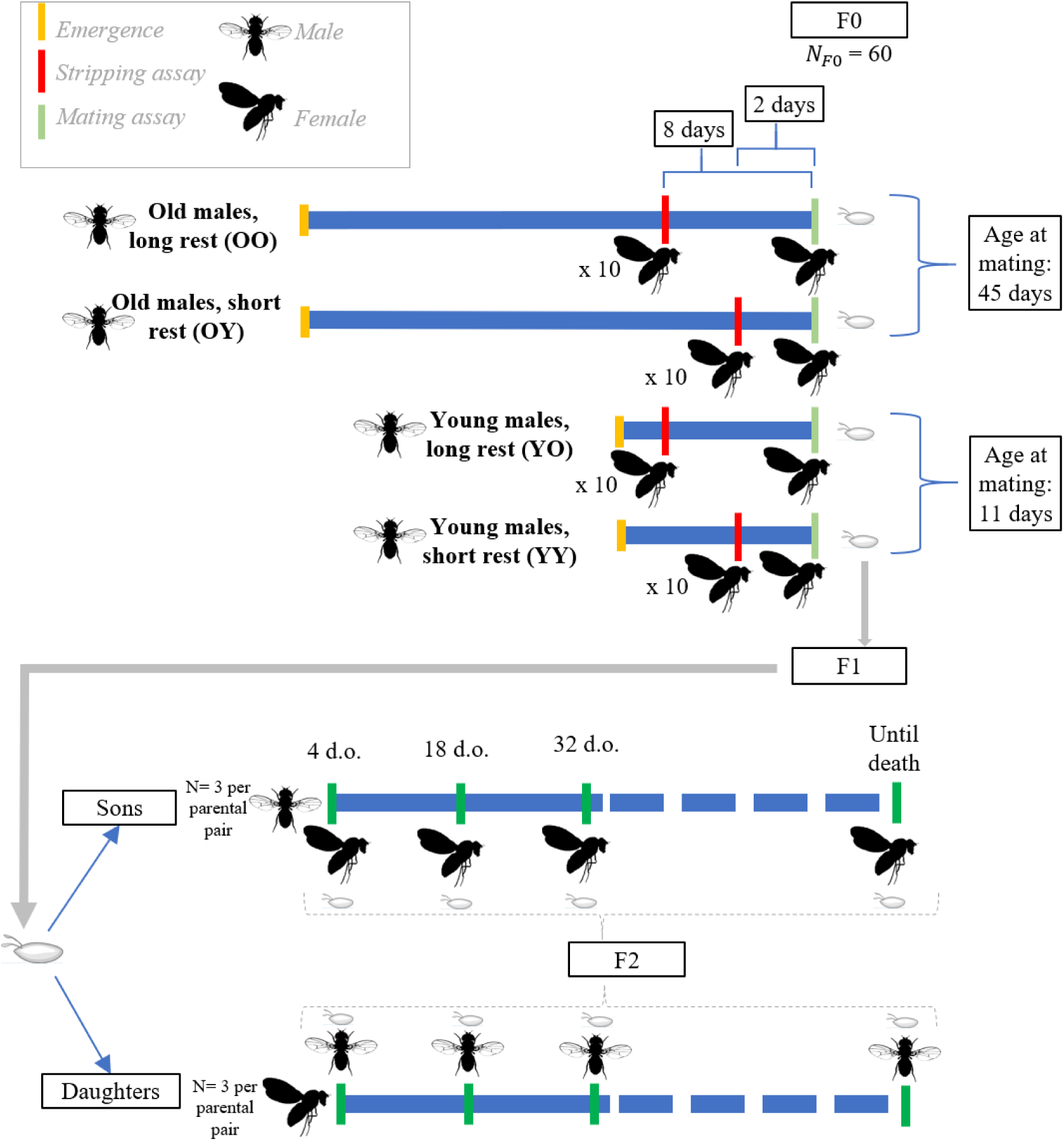
Experimental design to test how paternal age (Y: young, O: old) and paternal sperm storage duration/sexual rest (Y: short, O: long) affect lifetime reproduction and survival of sons and daughters. We first depleted old and young males of their ejaculates by keeping them with 10 females for 24 hours, then allowed them to replenish their ejaculate reserves for either two days (short sexual rest) or eight days (long sexual rest), and subsequently mated them to a single female. Three sons and three daughters produced from this mating (F1) were chosen haphazardly, and used for the F1 assays. Here, each experimental F1 offspring was mated to an individual of the opposite sex from the stock population once every two weeks, until death, and offspring reproductive output (number of eclosed F2 individuals) from eggs laid over 24 hours were counted. In total, each paternal treatment had 60 fathers. Images from PhyloPics by Thomas Hegna and Ramiro Morales Hojas (PD1.0 and CC01.0 licence).

## Experimental design

### F0 assays

#### Paternal treatments

We first reared experimental flies using a standard larval density method by placing ∼200 eggs obtained from our stock population cage, on 50 mL of food in 250-mL bottles (Clancy and Kennington, 2001). We then collected virgin “F0 males” using ice anaesthesia within 7 hours of eclosion from across ∼15 of these bottles. F0 males were kept in groups of 10 and haphazardly assigned to one of four paternal treatments. These four paternal treatments were: old fathers with sperm stored for long durations (OO), old fathers with sperm stored for short durations (OY), young fathers with sperm stored for long durations (YO), and young fathers with sperm stored for short durations (YY). To generate fathers with sperm of known storage durations, we manipulated the duration of sexual rest. For this, we “stripped” (i.e. depleted stored ejaculates) virgin young and old F0 males of their stored ejaculates (see “F0 stripping assay” below), and then allowed F0 males to replenish their ejaculates for a known duration, until they were mated to a single young virgin experimental female to obtain offspring (see F0 mating assay below). Specifically, F0 males assigned to the OO treatment were stripped when 37 days old and mated at 45 days old, OY males were stripped when 43 days old and mated at 45 days old, YO males were stripped when 3 days old and mated at 11 days old, while YY males were stripped when 9 days old and mated at 11 days old (Figure 1). This design gave us F0 males who were young (11 days old) or old (45 days old), and with ejaculates stored for short (2 days) or long (8 days) durations.

#### F0 stripping assay

To deplete (“strip”) experimental F0 males of their stored ejaculate reserves and create the four paternal treatments, we placed single old or young F0 males with 10 virgin young (3-4 days old) females (henceforth, “stripping females”), and allowed males to mate *ad-libitum* with these stripping females for 24 hours. Mating with 10 females over 24 hours is sufficient to deplete *D. melanogaster* males of their ejaculate reserves (Douglas et al, 2020; Hopkins et al, 2019; Linklater et al, 2007; Loyau et al, 2010; Macartney et al, 2021; also see Appendix 1). After 24 hours of being with the 10 stripping females, F0 males were separated, sexually rested for two or eight days depending on their treatment, and used subsequently for the “F0 mating assay”. Finally, we measured whether the F0 males used in the stripping assay produced offspring with the stripping females, to ensure the stripping assay’s effectiveness (see Appendix 1 for details).

#### F0 mating assay

After being with the 10 stripping females for 24 hours, all F0 males were transferred to new vials and kept individually. Only F0 males that produced offspring in the “F0 stripping assay”, were used in the “F0 mating assay” (see Appendix 1 for details). Next, old and young F0 males assigned to the short sperm storage treatment (OY and YY respectively) were mated once with a young virgin (3-4 days old) experimental female two days after the F0 stripping assay. Old and young F0 males in the long sperm storage treatment (OO and YO respectively) were mated once with a young virgin (3-4 days old) experimental female 8 days after the stripping assay. All experimental females (mothers) were obtained from standard stock cages using the standard larval density method described above. F0 males from all four treatments (henceforth called “fathers”) were presented with an experimental female on the same day, given a maximum of 5 hours to mate only once. Each mating event was observed, and mating latency and copulation duration recorded. Following mating, experimental females were left singly in the mating vial for 24h to enable oviposition. The resultant offspring (“F1”) produced by experimental females (mothers) and fathers, over 24 hours of eggs laying in mating vials, were used in the F1 assays (see below).

### F1 assays

The vials with eggs laid from experimental parents in the four treatments were checked every day for eclosing offspring (F1). Within seven hours of eclosion, three male (henceforth “sons”) and three female (henceforth “daughters”) offspring were haphazardly collected from each vial, into individual vials with a unique ID. These six offspring from each parental pair were then used for subsequent “F1 assays” (see below). The remaining eclosed offspring in each parental vial were frozen four days later and counted, to compare the reproductive output of fathers from the four treatments. Overall, we conducted F1 assays on 3 sons and 3 daughters from ∼60 fathers in each treatment.

Once every two weeks, surviving sons and daughters were moved to a new vial with a virgin young mate (3-4 days old) of the opposite sex for 24 hours and the resultant offspring (“F2”) were counted. This gave us data on the reproductive ageing patterns of both sons and daughters, from fathers in all four paternal treatments. We checked offspring survival every one to three days. Using an aspirator, sons were transferred to new food vials once a week, while daughters were transferred to new vials twice a week to reduce female mortality caused by larvae softening the food medium.

### Data analysis

#### General modelling approach

We used linear mixed-effects models (LMM) and generalised linear mixed-effects models (GLMM) to understand how paternal age and paternal sperm storage duration affected the reproductive output of fathers and their offspring (Table 1). We included an optimizer (“BFGS”) whenever GLMM models did not converge. We used Cox mixed-effects proportional-hazards models (coxme) to understand how paternal age and paternal sperm storage duration affected the age-dependent mortality of offspring. All analyses were done in R v3.5.2 (R Core team, 2012), using the packages *stats* (R Core team, 2012), *lme4* (Bates et al, 2015)*, glmmTMB* (Brooks et al, 2017)*, and coxme* (Therneau, 2015). All LMMs were checked for normality and homoscedasticity of residuals using the *stats* package, and GLMMs were checked for overdispersion whenever appropriate, using the *DHARMa* package (Hartig and Hartig, 2017). We analysed data on sons and daughters separately.

**Table 1:** Detailed model structure for each statistical model in our study. Model aim, dependent variables, fixed and random effects, and model type for the best-fitting model (determined using AIC comparisons) are reported. Proportion of variance in data explained by fixed effects reported as marginal R^2^.

| Generation | Model | Dependent variable | Fixed effects | Random effects | Model type | $R^2_{\text{marginal}}$ |
| --- | --- | --- | --- | --- | --- | --- |
| F0 (fathers) | Reproductive output | Number of offspring produced | Age*sperm storage + copulation duration + replicate |  | Zero-inflated negative binomial | 3.3% |
| F1 (offspring) | Reproductive ageing (daughters) | Number of offspring produced in a day of egg laying | Paternal age*paternal sperm storage*age + age^2 + lifespan + paternal fecundity + replicate | 1\|Paternal ID/daughter ID | Zero-inflated negative binomial | 38.8% |
|  | Reproductive ageing (sons) | Number of offspring produced in a day of egg laying | Paternal age*paternal sperm storage*age + lifespan + paternal fecundity + replicate | 1\|Paternal ID/son ID | Negative binomial | 2.6% |
|  | Actuarial ageing (daughters) | Lifespan | Paternal age*paternal sperm storage + paternal fecundity + replicate | 1\|Paternal ID | Cox-proportional hazards | 6.5% |
|  | Actuarial ageing (sons) | Lifespan | Paternal age*paternal sperm storage + paternal fecundity + replicate | 1\|Paternal ID | Cox-proportional hazards | 4.8% |

Marginal variance (R^2^) explained by fixed effects in our models, was calculated using the *sjPlot* (Ludecke, 2023) and *CoxR2* (You and Xu, 2020) packages. Post-hoc pairwise comparisons whenever conducted, were done using Hedges’ g in the *effectsize* (R Core team, 2012) package (α = 0.05 for significance tests of effect size). Model comparisons were done using AIC function in the *stats* package.

Unless mentioned otherwise, we started with a “full model” that included two-way interactions between paternal sperm storage duration and paternal age. For models on reproductive ageing in offspring, our full model additionally included offspring age in a three-way interaction with paternal age and sperm storage. These full models were used to interpret the highest order interactions only. To then interpret lower order interactions or main-effects whenever higher order-interactions were non-significant, we fitted models with the highest level of interaction removed (following Engqvist, 2005). Main effects indicated independent effects of variables when averaged across the effects of other variables.

### F0 assays

We analysed how paternal age and sperm storage duration affected the reproductive output of fathers. We modelled the number of offspring produced by fathers as our dependent variable, with zero-inflated negative binomial error distribution, because this distribution fit data better than Poisson (ΔAIC = 6624, ΔDF = 2) or zero-inflated Poisson (ΔAIC = 1283, ΔDF = 1) error distributions. We modelled paternal age, sperm storage duration, their interaction, copulation duration, and replicate as fixed effects. Copulation duration was included to account for males who copulate for longer durations transferring more sperm to females, thus producing more offspring.

### F1 assays

#### Reproductive ageing (Table 1)

To understand the effects of paternal treatment on age-dependent reproductive output of daughters, we built a GLMM with zero-inflated negative binomial error distribution. This model was a better fit to the data than one with Poisson (ΔAIC = 20967, ΔDF = 3) or zero-inflated Poisson (ΔAIC = 3151, ΔDF = 1) error distribution. We modelled the number of offspring produced over 24 hours by daughters, measured once every two weeks from birth to death, as the dependent variable. We included paternal age, paternal sperm storage duration, the age of daughters, their three-way interaction, and replicate as fixed effects. In the same model, we included a quadratic term for daughter’s age (which significantly improved model fit compared to only a linear term, P < 0.001), because reproductive ageing patterns are often curvilinear (Jones et al, 2014; Sanghvi et al, 2024). Effects of paternal age on offspring lifetime reproduction might be biased, if demographic processes such as selective disappearance cause non-random death of offspring over time. In our model, we thus additionally included the lifespan of daughters as a fixed effect, to account for selective disappearance (Bouwhuis et al, 2009; Sanghvi et al, 2022). We further included the number of offspring that fathers produced as a fixed effect, to investigate whether fathers compensate for lower quality offspring by producing more offspring. We modelled daughter ID nested within paternal ID as random effects.

To understand the effects of paternal treatment on age-dependent reproductive output of sons, we built a GLMM with a negative binomial error distribution. This model was a better fit to the data than one with Poisson (ΔAIC = 13177, ΔDF = 1) or zero inflated Poisson (ΔAIC = 4211, ΔDF = 0) error distributions. The fixed and random effects in the model for reproductive output of sons were identical to those in our model for reproductive output of daughters (as described above), except for one difference. Specifically, for sons, we modelled only a linear term for the age at reproduction of sons, because a quadratic term did not improve model fit (ΔAIC = 2, ΔDF = 1, P = 0.766 with L.R.T. under a Chi-sq. distribution).

#### Actuarial ageing (Table 1)

We used Cox- mixed-effects proportional hazards models to investigate the age-dependent mortality risk to sons and daughters separately. However, our models for both sons and daughters were identically structured. We modelled the lifespan of offspring as the dependent variable. Paternal age, paternal sperm storage duration, their two-way interaction, and replicate were included as fixed effects. The number of offspring sired by fathers was modelled as a fixed effect, to test whether fathers compensate for low survival of offspring by producing more offspring. We modelled paternal ID as a random effect.

## Results

### F0 assays

We found no interaction between paternal age and sperm storage duration influencing the number of offspring produced by fathers (z = 0.343, P = 0.731, Figure 2B, Table S2).

**Figure 2:**
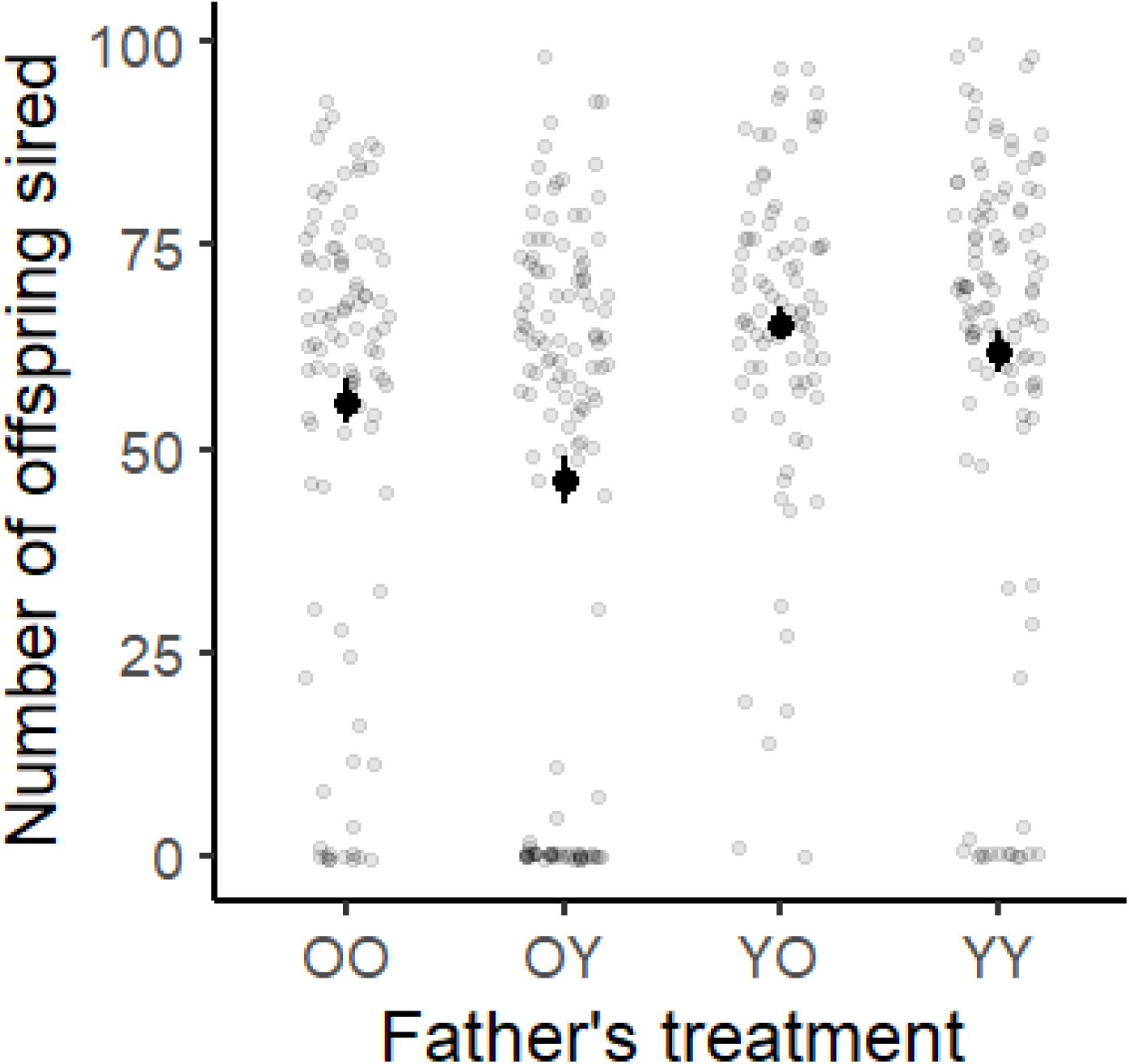
Effects of age and sperm storage treatment, on the number of offspring sired by fathers across the four paternal treatments: old age, long sperm storage (OO); old age, short sperm storage (OY); young age, long sperm storage (YO); young age, short sperm storage (YY). Means and SE shown along with the raw data points.

However, paternal age (z = 2.130, P = 0.033), but not sperm storage duration (z = 0.810, P = 0.419, Figure 2B), independently affected the number of offspring fathers produced, with young fathers producing more offspring than old fathers.

### F1 (offspring) assays

#### Reproductive ageing

The three- or two- way interaction of paternal age, paternal sperm storage duration, and the age of daughters, had no significant effect on the number of offspring produced by daughters (Figure 3, Table S3). Furthermore, paternal age (z = -1.14, P = 0.254) or paternal sperm storage duration (z = 1.43, P = 0.153) did not significantly influence the reproductive output of daughters (Table S3). However, daughters produced fewer offspring when old than when young (daughter age as quadratic: z = -3.410, P = 0.001; as linear: z = -6.050, P < 0.001).

**Figure 3:**
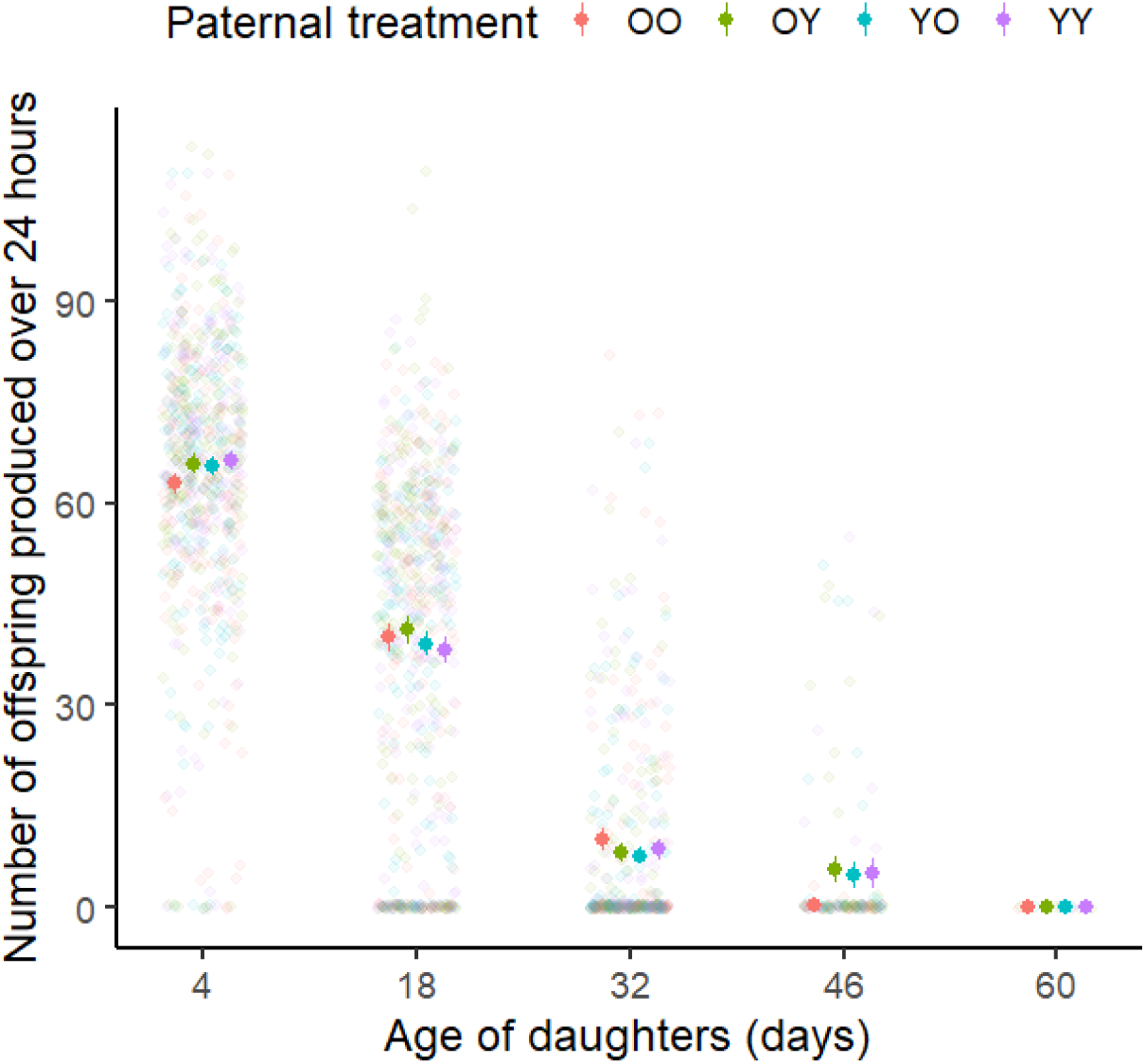
No effect of paternal age or paternal sperm storage duration on the number of offspring produced by daughters. OO: Old paternal age, long paternal sperm storage; OY: Old paternal age, short paternal sperm storage; YO: young paternal age, long paternal sperm storage; YY: young paternal age, short paternal sperm storage. Means and SE shown along with raw data. Each light dot represents measurements on a single daughter for a given age.

Daughters that lived longer also produced more offspring at a given age, than daughters that lived shorter lives (daughter’s lifespan: z = 2.710, P = 0.007, Figure 4A), a result consistent with selective disappearance. Paternal reproductive output did not influence the reproductive output of daughters (z = -0.020, P = 0.984, Figure S3A).

**Figure 4:**
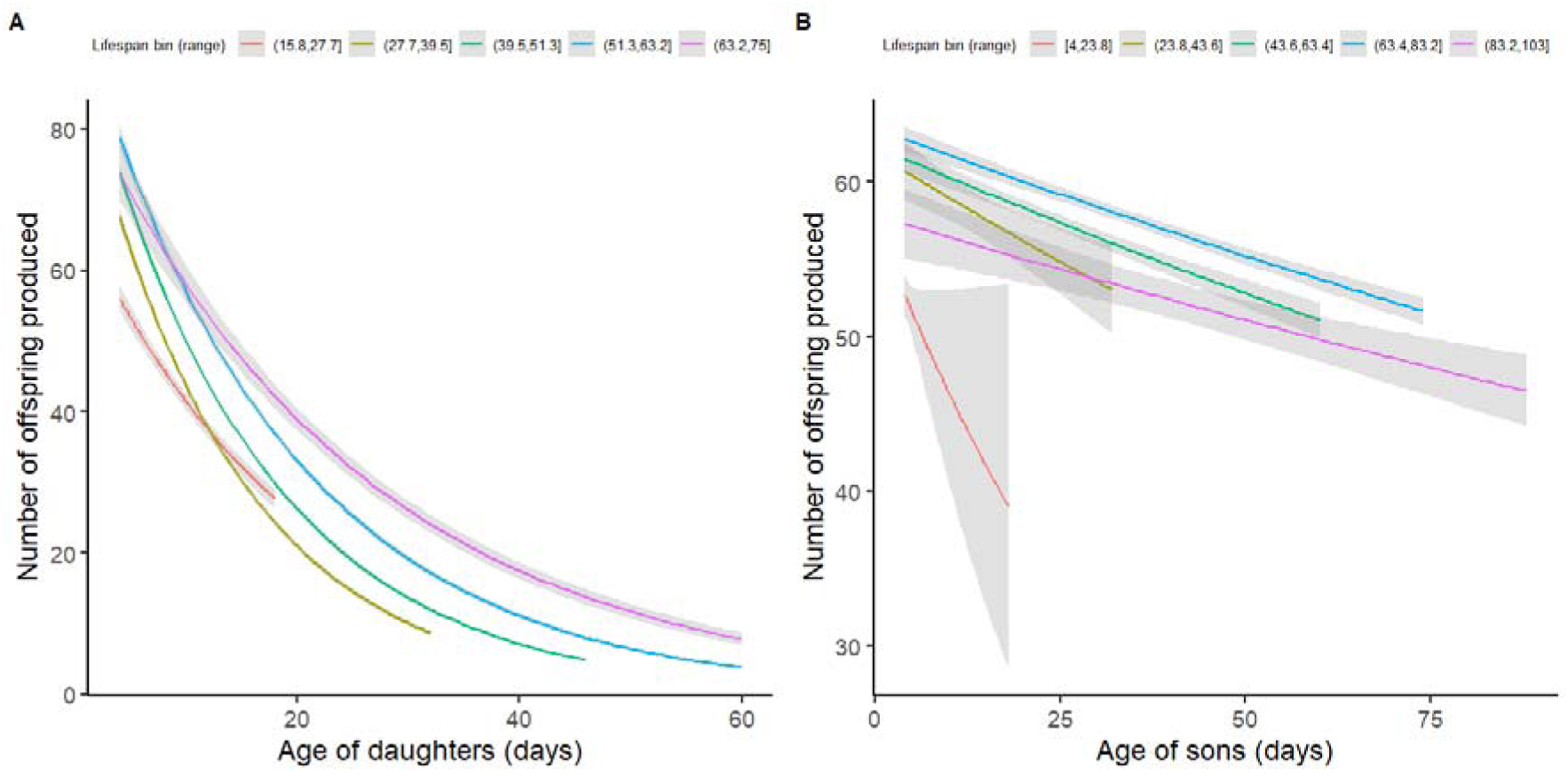
A- Daughters who lived longer consistently produced more offspring throughout life, than daughters who lived shorter lives, suggesting selective disappearance. B- We found no significant evidence for selective disappearance in sons. Data binned within 5 lifespan ranges. Curves plotted as glm-poisson. Shaded areas represent 95% C.I.

We found no significant effect of three-way or two-way interactions between paternal age, paternal sperm storage duration, and the age of sons, to affect the number of offspring produced by sons (Figure S4, Table S4). Similarly, paternal age did not have a significant effect (z = 0.160, P = 0.875). However, we found a marginally significant effect of paternal sperm storage duration (z = 2.030, P = 0.043, Figure 5A) on the number of offspring produced by sons. Overall, sons born to fathers who stored sperm for eight days (i.e. long sperm storage) had a 4.2% lower reproductive output than sons born to fathers who stored sperm for two days. Post-hoc tests revealed that the magnitude of this difference was greatest in the early life of sons (Hedge’s g at 4 days old: -0.22, 6.5% difference; Figure 5A, 5B). The number of offspring sired by fathers was positively correlated with the number of offspring sired by their sons (z = 2.930, P = 0.003, Figure S5). However, unlike in daughters, we did not find significant evidence for selective disappearance in sons (son’s lifespan: z = 1.660, P = 0.097; Figure 4B). When averaged across other variables, sons produced fewer offspring with advancing age (z = -4.43, P < 0.001).

**Figure 5:**
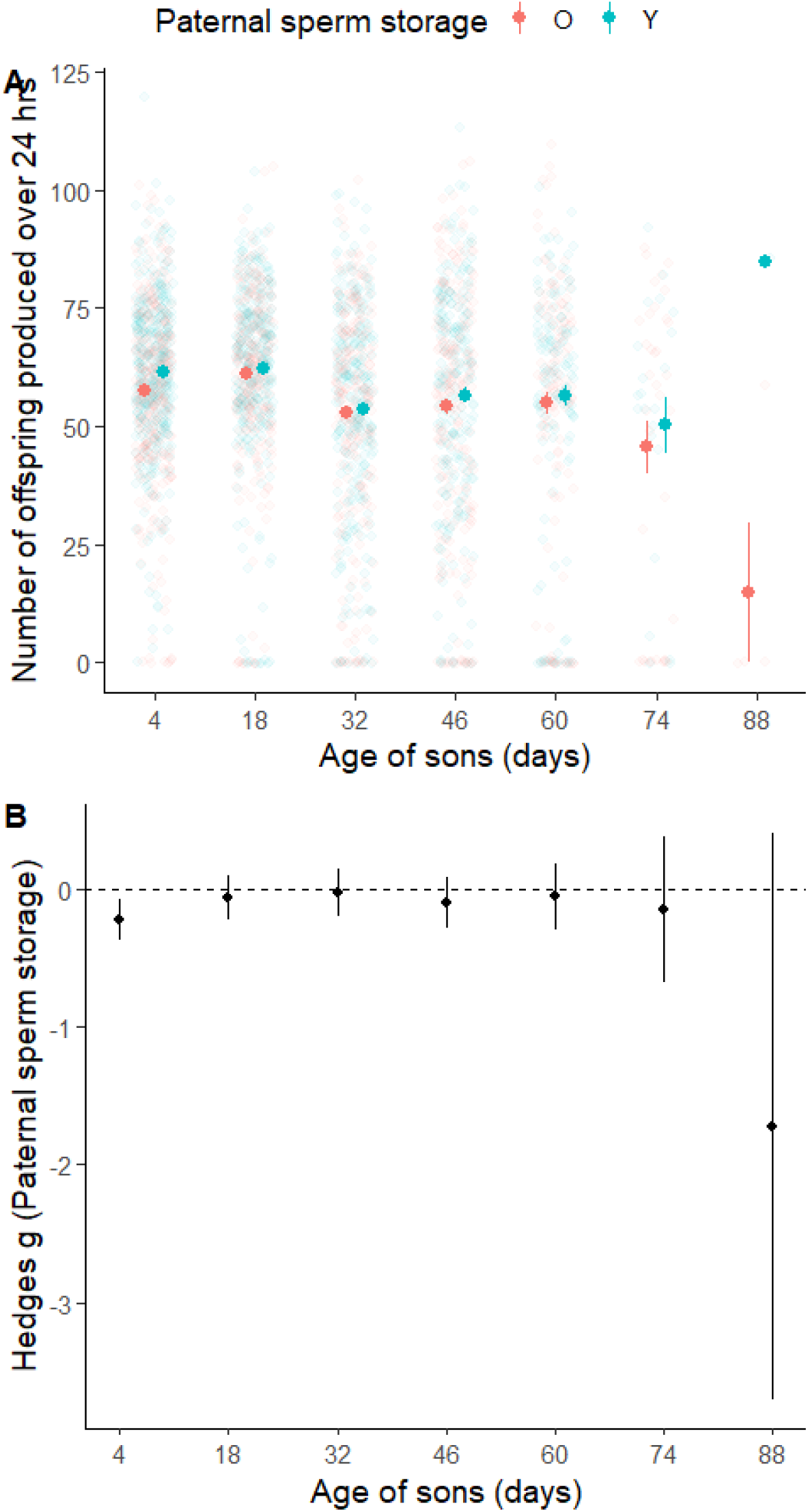
A- Effect of paternal sperm storage duration on the number of offspring produced by sons, when averaged across effects of other variables. Means and SE shown along with raw data. O: long paternal sperm storage, Y: short paternal sperm storage. Each light dot represents measurements on a single son at a given age. B- Effect of paternal sperm storage duration on reproductive success of sons is significant when sons are 4 days old. Effect sizes (Hedges; g) used for comparisons, and significance tests based on whether the 95% C.I. overlaps with zero or not. Negative effect sizes indicate that sons from fathers with long sperm storage duration, have lower reproductive output than sons from fathers with short sperm storage. Means and C.I. shown.

#### Actuarial ageing

Paternal age, paternal sperm storage duration, or their two-way interaction, did not significantly influence age-dependent mortality risk of daughters (Figure 6A, Table S5) or sons (Table S6). However, the number of offspring sired by fathers was significantly correlated with the mortality risk of their sons (z = -2.260, P = 0.024, Figure S5), but not daughters (z = 0.500, P = 0.610, Figure S5). Specifically, fathers who produced more offspring conceived sons that had higher rates of survival.

**Figure 6:**
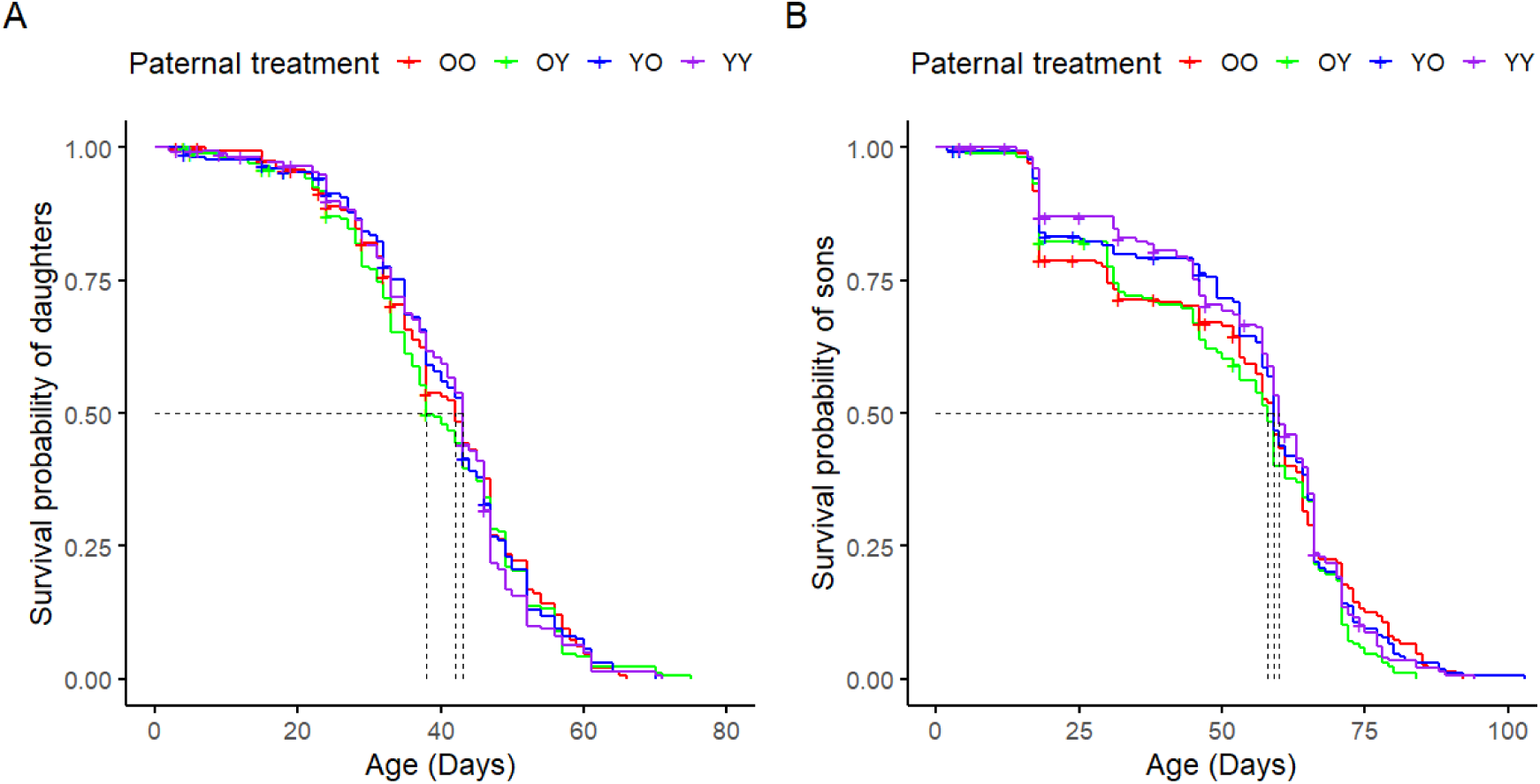
No significant effect of paternal age or paternal sperm storage duration on the age-dependent survival probability of A- daughters, or B- sons. Dotted lines show age at median survival probability. OO: old paternal age, long paternal sperm storage; OY: old paternal age, short paternal sperm storage; YO: young paternal age, long paternal sperm storage; YY: young paternal age, short paternal sperm storage. “+” shows censoring (lost, accidentally killed, or mislabelled).

## Discussion

Paternal age effects can have far-reaching ramifications for organismal evolution. However, determining the causal effect of paternal age on offspring phenotype and fitness requires experimental control of potential confounds, which is challenging. We investigated how paternal age and paternal sperm storage duration affect the reproductive output of fathers, and the reproductive and actuarial ageing of their offspring. We found older fathers to have lower reproductive output, suggesting reproductive senescence, but found no significant evidence for paternal age effects on offspring (H1). Additionally, we did not find evidence for paternal sperm storage duration interacting with paternal age, to influence offspring phenotypes (H2). Instead, we found weak evidence for an independent effect of paternal sperm storage duration on the reproductive output of sons. Furthermore, the lack of paternal age effects was consistently observed irrespective of offspring age (H3). We also found no evidence for paternal age to affect sons more than daughters, but observed this pattern for paternal sperm storage duration (H4). Lastly, there were no trade-offs between offspring quantity versus quality produced by fathers. Instead, fathers that produced more offspring also produced higher quality sons (H5).

We found no significant effect of paternal age on offspring, when separated from effects of paternal sperm storage duration (H1). Paternal age effects have been widely reported across taxa (e.g. Crow, 2003; Monaghan et al, 2020; Schroeder et al, 2015; Vuarin et al, 2019), including in humans (Chang and Robaire, 2022), with offspring from old fathers having lower fitness than offspring from young fathers (reviewed in Monaghan and Metcalfe, 2019). However, most studies on paternal age effects do not control for duration of sperm storage. The lack of an effect in our study might suggest some of the previous reports misattributing deleterious effects of paternal sperm storage to paternal age (reviewed in Pizzari et al, 2008, Sanghvi et al, 2024). In fruit flies for example, experimental males are often unmated until they are used to sire progeny (e.g. Aguilar et al, 2023; Mossman et al, 2019; Nystrand and Dowling, 2014; Price and Hansen, 1998; Priest et al, 2002). Male fruit flies are characterised by spermatogenesis throughout their adult life (Bjork et al, 2007; Sepil et al, 2020) and low rates of sperm loss (Demarco et al, 2014; Santos et al, 2023). These might lead to the accumulation of deteriorating sperm within the male reproductive tract, such that old virgin fathers have more deteriorated sperm than young virgin fathers. Only few studies have attempted to separate paternal age versus sperm storage effects, albeit in other species. For example, Gasparini et al (2019) report independent effects of both, paternal age and sperm storage duration, on paternal fertility in guppies; Jones et al (2004) show interactive effects between paternal age and sperm storage duration on paternal fertilisation success, in hide beetles; Vega-Trejo et al (2019) show no effect of paternal age or sperm storage duration on paternal sperm traits, in mosquitofish; while Meunier et al (2022) show effects of paternal age but not of sperm storage duration on sperm traits and offspring survival, in bustards. None of these studies however, measured offspring lifetime reproductive success (the key metric of evolutionary fitness), as we were able to do in the present study.

The lack of a paternal age effect could also reflect deleterious effects being balanced by beneficial effects of having old fathers (Sanghvi, Pizzari et al, 2024). For instance, fathers who have survived to older ages might represent a biased subset of fathers with alleles that confer longer lifespans (due to viability selection: Brooks and Kemp, 2001; Hansen and Price, 1995; Johnson and Gemmell, 2012; Kokko, 1998; Sanghvi, Pizzari et al, 2024).

Similarly, fathers of poor reproductive value could selectively disappear with age (Bouwhuis et al, 2009; Hamalainen et al, 2014; Sanghvi et al, 2022), leading to old surviving fathers having alleles for higher reproductive output. These processes (selective disappearance and viability selection) might have underestimated paternal age effects in our study, due to fathers being sampled-cross sectionally rather than the same fathers being sampled longitudinally.

The lack of a paternal age effect in our study could also occur due to female-driven processes. For example, female fruit flies might bias fertilisation toward good quality sperm via cryptic female choice (e.g Hadlow et al, 2023; reviewed in Firman et al, 2017; Sanghvi et al, 2024; Vuarin et al, 2019).

We did not find evidence that old fathers who stored sperm for long durations, produce lower quality offspring than other treatments (H2). Old males are hypothesized to have poorer DNA repair machinery than young males (Chen et al, 2023; Gorbunova et al, 2007; Selvaratnam et al, 2015; Weirich-Schwaiger et al, 1994), which could lead to old fathers being worse at repairing sperm damage than young fathers (Pizzari et al, 2008). The lack of an interaction between paternal age and sperm storage duration in our study could be due to sperm storage for eight days not being sufficient to increase mutation load in paternal sperm, or old fathers being able to repair DNA damage in mature sperm, well enough to ameliorate deleterious effects of sperm storage.

We found weak evidence for deleterious effects of paternal sperm storage duration on the early-life reproductive output of sons. Several studies have demonstrated that storage of mature spermatozoa can deteriorate sperm quality (Agarwal et al, 2016; Brindle et al, 2023; Cattelan and Gasparini, 2021; Comar et al, 2017; Gasparini et al, 2014, 2019; Hettyey et al, 2012; Levitas et al, 2005; Radhakrishnan and Fedorka, 2011), and lead to offspring having lower fitness (Gasparini et al, 2017; Wagner et al, 2004; White et al, 2008). The likely mechanism causing such deleterious effects is sperm storage increasing oxidative stress in sperm and causing increased sperm DNA fragmentation (Barbagallo et al, 2022; Wetzker et al, 2024). Other studies however, have not obtained evidence for such deleterious effects (Firman et al, 2015; Hotzy et al, 2020; Meunier et al, 2022; Vega-Trejo et al, 2019). There are several explanations for only a weak effect of paternal sperm storage duration on offspring being recorded in our study. First, sperm could be continuously re-absorbed or lost in fathers, leading to low levels of sperm damage despite long durations of sexual rest (Pizzari et al, 2008; Reinhardt and Siva-Jothy, 2005; Reinhardt, 2007). Second, a weak effect could indicate sperm not being stratified in males (Reinhardt, 2007), causing fresh and stored sperm to be mixed in paternal ejaculates. Third, it is possible that not all the sperm stored in male reserves were ejaculated by F0 males in our stripping assay (Sanghvi, Shandilya et al, 2024), leading to less effective sperm storage treatments. Fourth, if sperm haploid genomes are expressed in fruit flies, selection on sperm haplotypes could lead to the death of poor-quality sperm in fathers who stored sperm for long durations (Alavioon et al, 2017; Immler, 2019; Otto et al, 2015). This selective death could buffer deleterious effects of paternal sperm storage duration on offspring. Fifth, females could be actively ejecting sperm stored for longer durations in males, thus buffering the effects of sperm storage (Reinhardt and Siva-Jothy, 2005; Snook and Hosken, 2004). In our study, the effect of sperm storage on the reproductive output of sons was significant, despite its magnitude being small. This statistical significance could thus be an artefact of having a large sample size (of ∼180 sons per treatment) rather than a true biological pattern. We only manipulated sperm storage duration in fathers, but future studies can investigate whether sperm storage in mothers (reviewed in Orr and Brennan, 2015) also influence offspring phenotypes.

We predicted (H3) that deleterious effects of old paternal age would be more severe for old than young offspring (Brengdahl et al, 2023; Moorad and Promislow, 2008; reviewed in Monaghan et al, 2020). Such an interaction could occur if senescence in offspring exacerbated effects of a higher mutation load inherited from old parents (Chen et al, 2023; Girard et al, 2016; Kong et al, 2012; Yatsenko and Turek, 2018). Studies on fruit flies show that male germlines accumulate mutations with age (Garcia et al, 2010; Wang et al, 2022), that age-specific mutation accumulation can lead to accelerated senescence (Brengdahl et al, 2020; Yampolsky et al, 2000), and that old fathers have a higher germline mutation load than young fathers (Witt et al, 2023). A lack of support for the predicted interactive effect in our study might indicate that inherited mutations in fruit flies are mostly selectively neutral (reviewed in de Jong et al, 2023) in their fitness effects. Selective disappearance could have also masked such an interactive effect in daughters, where daughters of worse reproductive quality selectively died with age. Overall, we encourage future studies to not only quantify age-dependent mutation rates and epigenetic changes in old paternal sperm (e.g. Oakes et a, 2003; Suvorov et al, 2020), but also investigate their inter-generational phenotypic effects.

We did not find paternal age to have sex-specific effects on offspring (H4), likely due to an overall absence of paternal age effects in our study. This lack of evidence also suggests that mechanisms of inheritance causing sex-specific paternal age effects, might be absent in fruit flies. Lastly, we did not find evidence that fathers who produced more offspring also produced offspring of poorer quality due to a trade-off between these traits (H5). Instead, paternal fecundity co-varied positively with the survival and reproductive output of their sons. These results might be explained by heterogeneity in paternal condition, such that fathers who can acquire more resources are able to invest in both, offspring production (e.g. via higher ejaculate production) and offspring quality (e.g. via sperm repair) without trade-offs (Reznick et al, 2000; Roff and Fairbairn, 2007). Due to these covariances being apparent only in sons but not in daughters, these results might alternatively be explained by sex-specific heritability of fitness-components (Calsbeek et al, 2015; Connallon, 2012; Weiss et al, 2006) in fruit flies.

## Conclusions

Our study challenges the commonly held prediction that older fathers produce lower quality offspring, and we find no evidence for deleterious paternal age effects. The absence of a paternal age effect persists irrespective of the age of sons and daughters, or the duration of paternal sperm storage. These results call for a re-evaluation of the causes and consequences of paternal age effects, under a framework that incorporates selective disappearance, condition dependence, life-history trade-offs, and variable paternal mating rates. We emphasize that post-meiotic sperm damage during sperm storage/sexual rest might be an under-appreciated modulator of offspring phenotypic variation, especially in species where low male mating rates and long-term female sperm storage are commonplace. The interactive influence of male age and sexual rest on male reproductive output and his offspring’s phenotypes, might be crucial in modulating female sperm ejection (Wagner et al, 2004), mate choice (Johnson and Gemmell, 2012), polyandry, and last-male sperm precedence (Snook and Hosken, 2014). Overall, we highlight the importance of simultaneously understanding various mechanisms by which advancing organismal age might influence offspring directly (e.g. sex-specific inheritance, mutation accumulation) or indirectly (e.g. sperm storage, selective disappearance, trade-offs, condition dependence).

## Supplementary information statement

Supplementary information is provided along with this manuscript

## Data availability statement

Data and associated code for this study are available at OSF under an anonymous folder: https://osf.io/gq9j2/?view_only=362a13e2bd6f4b958e5fba66e7c75a2e

