## Supplementary information for "No evidence for paternal age effects on sons or daughters, when accounting for paternal sperm storage"

Appendix 1: F0 stripping assay

To create our four paternal treatments, we used old and young experimental F0 males, and kept each male with 10 females in a vial for 24 hours, to deplete males of their ejaculate. This was done so that after depletion of ejaculates, males would produce sperm of a known maximum post-meitoic storage duration. After 24 hours of being with a female, we kept males in individual vials allowing either two or eight days of sexual rest. For all F0 males used in the stripping assay (and eventually in the paternal treatments), we ensured that the 10 females associated with him, collectively, produced at least one offspring. This was done to ensure that all the F0 males that were to be used in subsequent parts of the experiment in the four paternal treatments, were fertile.

In addition to this stripping assay, we tested whether F0 males had mated with the females they were kept with, in the stripping assay. For this, we chose a random subset of F0 males (see Table S1 for sample sizes), and collected each of the 10 females he was with, on individual vials. For each male, we recorded the number of associated stripping females he was with, which produced an offspring (Figure S2A). On average, males from all four paternal treatments produced offspring with >9 out of 10 females (Figure S2A). This “stripping fertility” assay, was done using eggs laid by females within 24 hours of mating.

Next, we chose another subset of F0 males (see Table S1 for sample sizes) and moved each of the ten females he was with into individual vials for 24 hours for egg laying. This “stripping counts” assay was done three days after the F0 “stripping fertility” assay. We then counted all the offspring that emerged from laid eggs. This “stripping counts” assay was done to ensure that males did not maintain their level of offspring production across all 10 females, and instead showed a large variation in how many offspring they produced with each female (Figure S2B, S2C), with many females producing very few offspring. Such a pattern in female fecundity would be indicative of males having lower numbers of remaining sperm when mating with each successive female through a mating sequence (Douglas et al, 2020; Hopkins et al, 2019; Linklater et al, 2007; Loyau et al, 2010; Macartney et al, 2021; Sirot et al, 2009; Figure S2B, S2C). On the other hand, maintenance of offspring production across females would indicate that males still had a lot of sperm stored, making our stripping assay ineffective in creating the two sperm storage duration treatments.

To further ensure that ten matings were sufficient to deplete males of their sperm reserves, we used a transgenic line of *Drosophila melanogaster* males that express green fluorescent protein in their sperm heads at the *Mst35Ba* and *Mst35Bb* loci (Manier et al, 2010). We dissected two virgin males who were 6-days old and compared the numbers of sperm in their seminal vesicles to two, 6-day old males who had been with 10 females in a vial for 24 hours. Visual comparison indicated many sperm in the virgin males’ seminal vesicles but almost no sperm in the mated males’ seminal vesicles (Images 1a and 1b respectively).

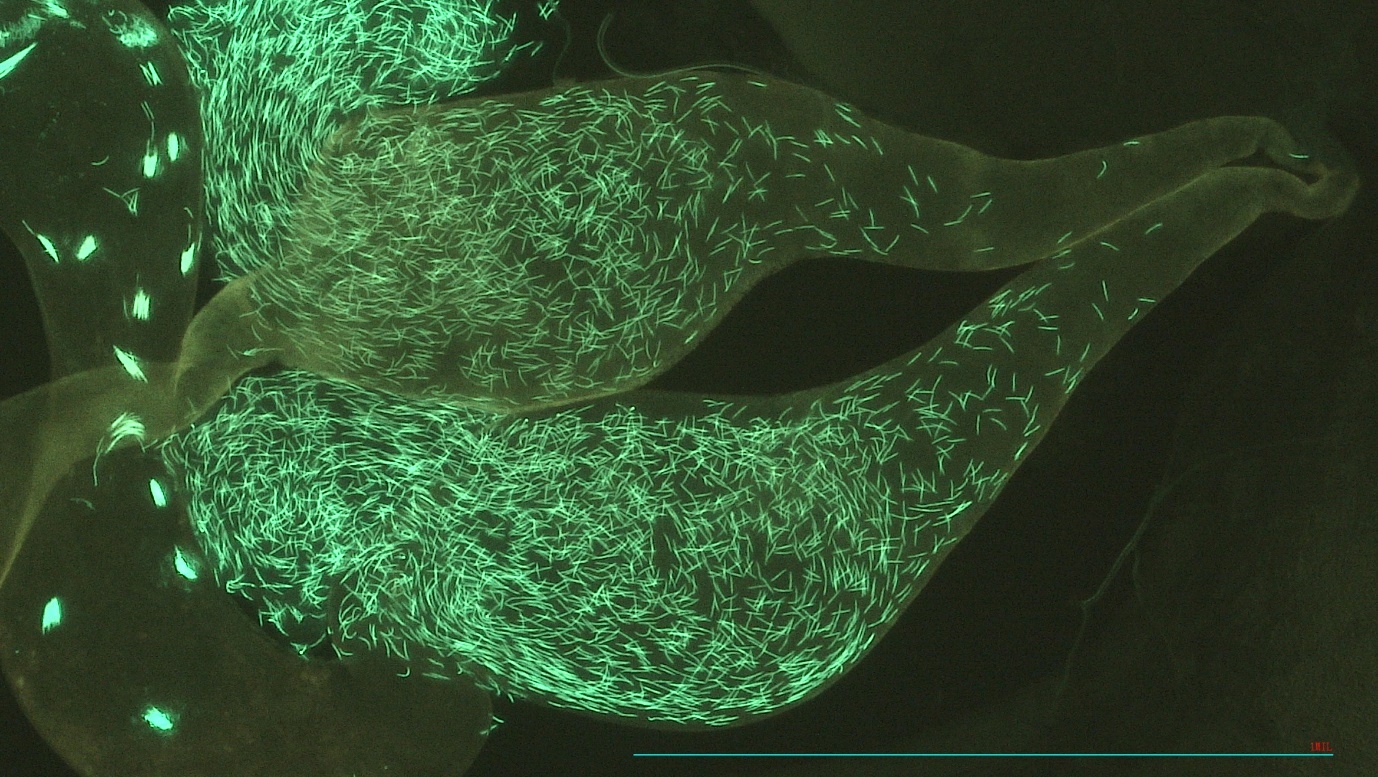

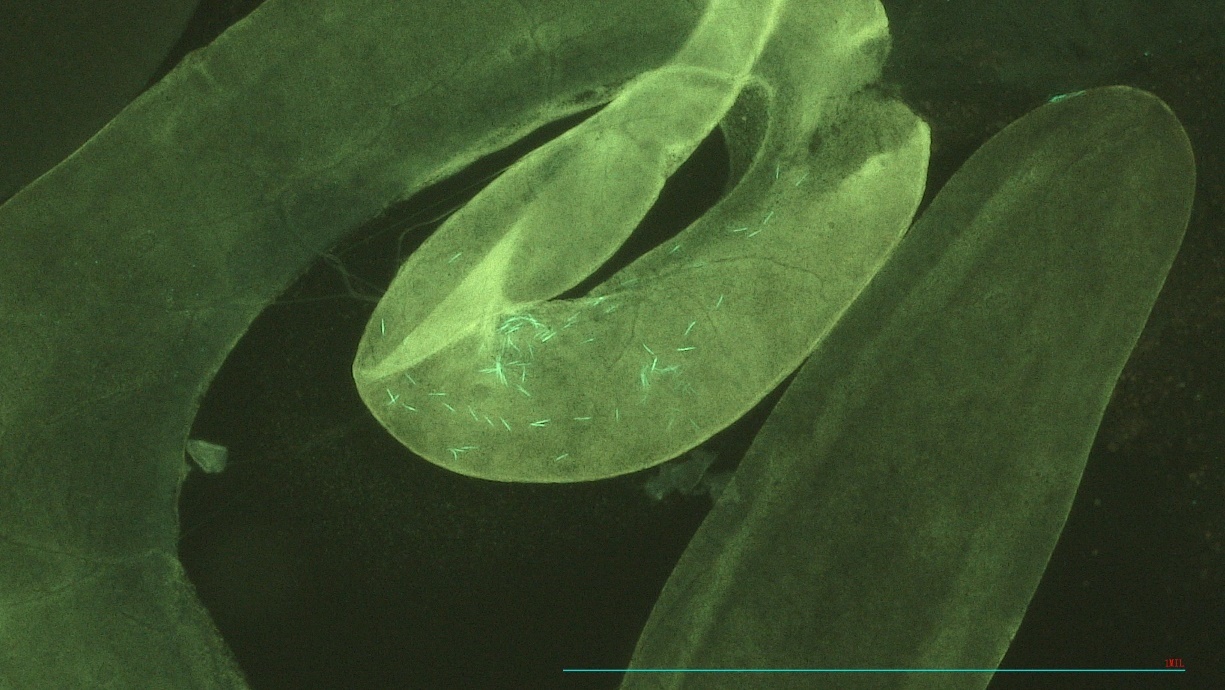

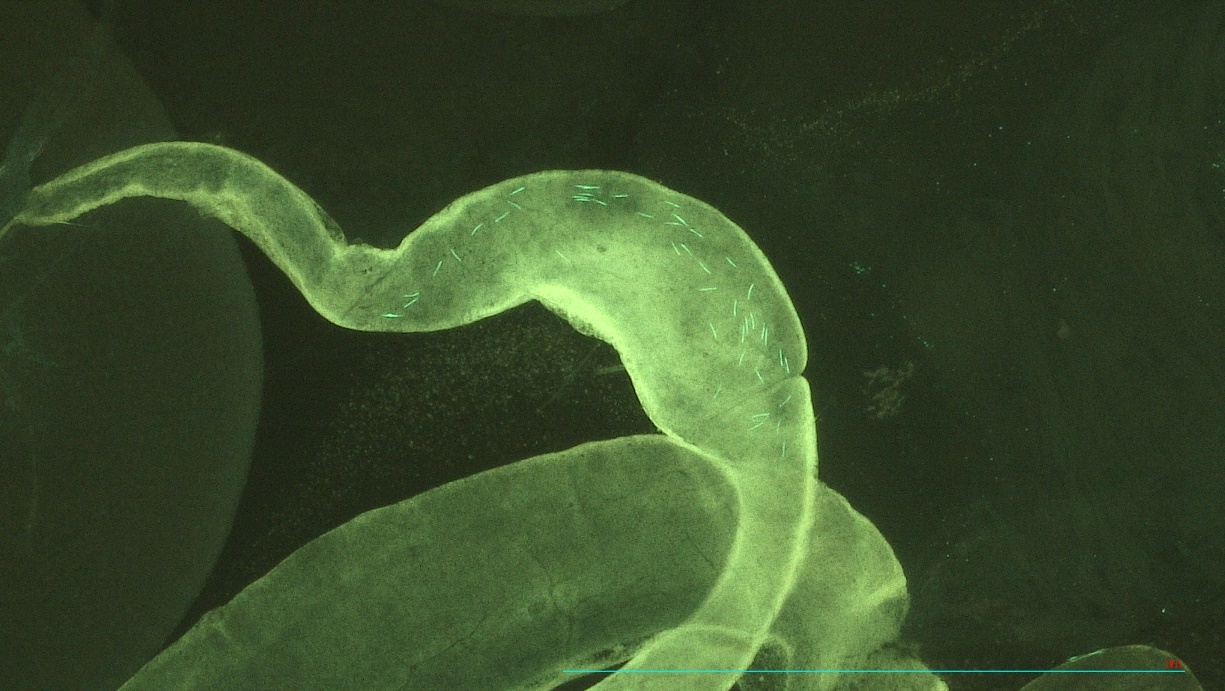

B

A

Image 1: A. GFP tagged sperm heads (bright green lines) in the seminal vesicles of virgin males that were six days old. B. GFP tagged sperm heads (bright green lines) in the seminal vesicles of six-day old males, who were kept in a vial with 10 females for 24 hours. Images taken using a Nikon Eclipse50i fluorescence microscope (magnification = 10x objective, 10x eyepiece, wavelength = 480nm) with a chromix HD camera, under UV light from a CoolLED pe300 light source

Appendix 2: Supplementary figures

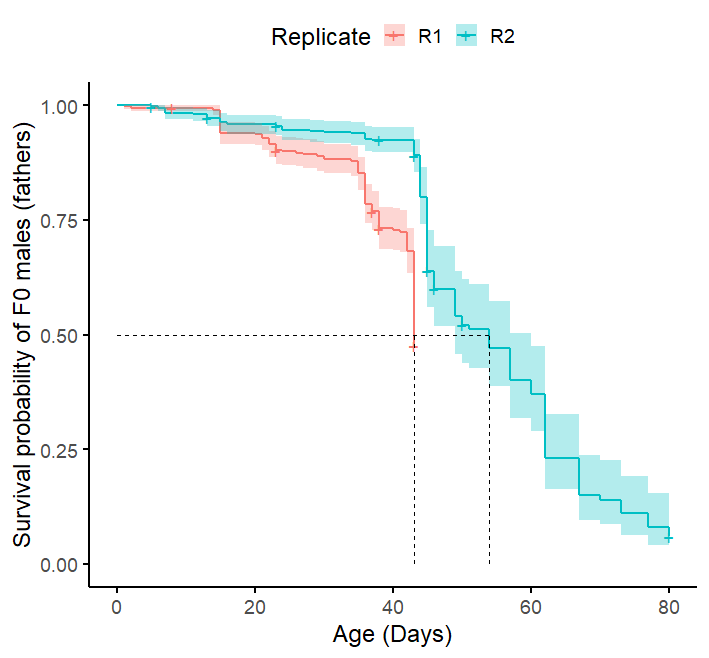

Figure S1: Survival probabilities of F0 males (kept in single-sex groups of 10) from replicates R1 and R2 used in our experiment. Initial sample size = ~370 for both replicates. Median lifespan is ~ 45 days. “+” shows censoring, shaded areas show 95% C.I. Lifespan assays for F0 males were only conducted in R1 and R2 due to logistical reasons.

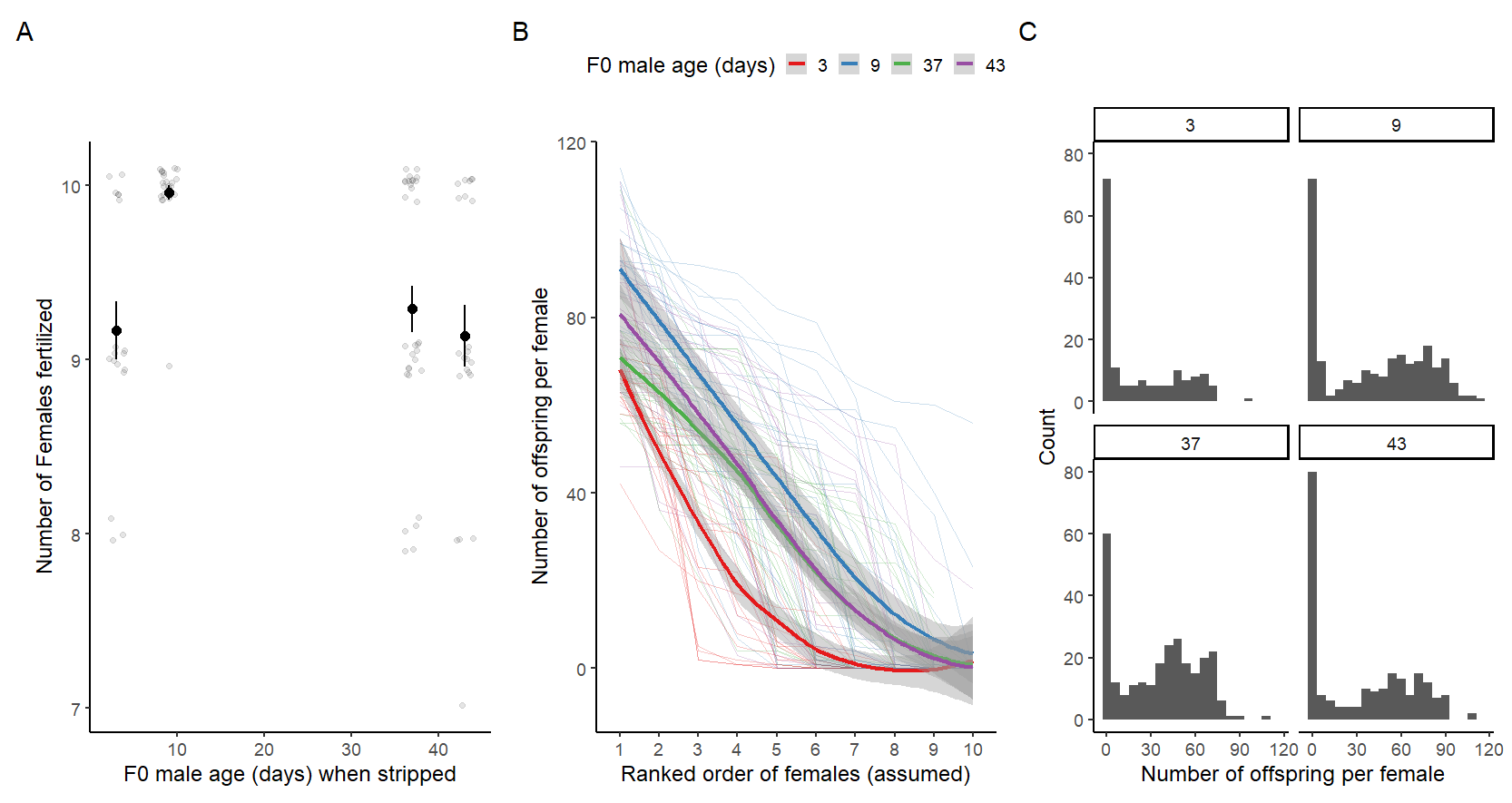

Figure S2: **A-** Number of females fertilized by F0 males (fathers) in the stripping assay (>9 out of 10 females on average; means and SE along with raw data presented). **B-** When all 10 stripping females from a subset of males were put on individual vials, and the number of offspring they produced counted, there was a large amount of variation in the number of offspring across females. For each F0 male assayed, we ranked females he was kept with, ascendingly from one to ten, based on the number of offspring they produced. This ranking was assumed to be the order in which females mated to a male, such that males produced fewer offspring with each successive female in a mating sequence. Dark lines show means of each F0 age group, light lines show individual males. Shaded areas show 95% C.I. **C-** Distribution of the number of offspring produced by each of the 10 stripping females a male was kept with, in the stripping-counts assay. F0 males of 3, 9, 37, and 43 days of age were assigned to YO, YY, OO, OY paternal treatments, respectively.

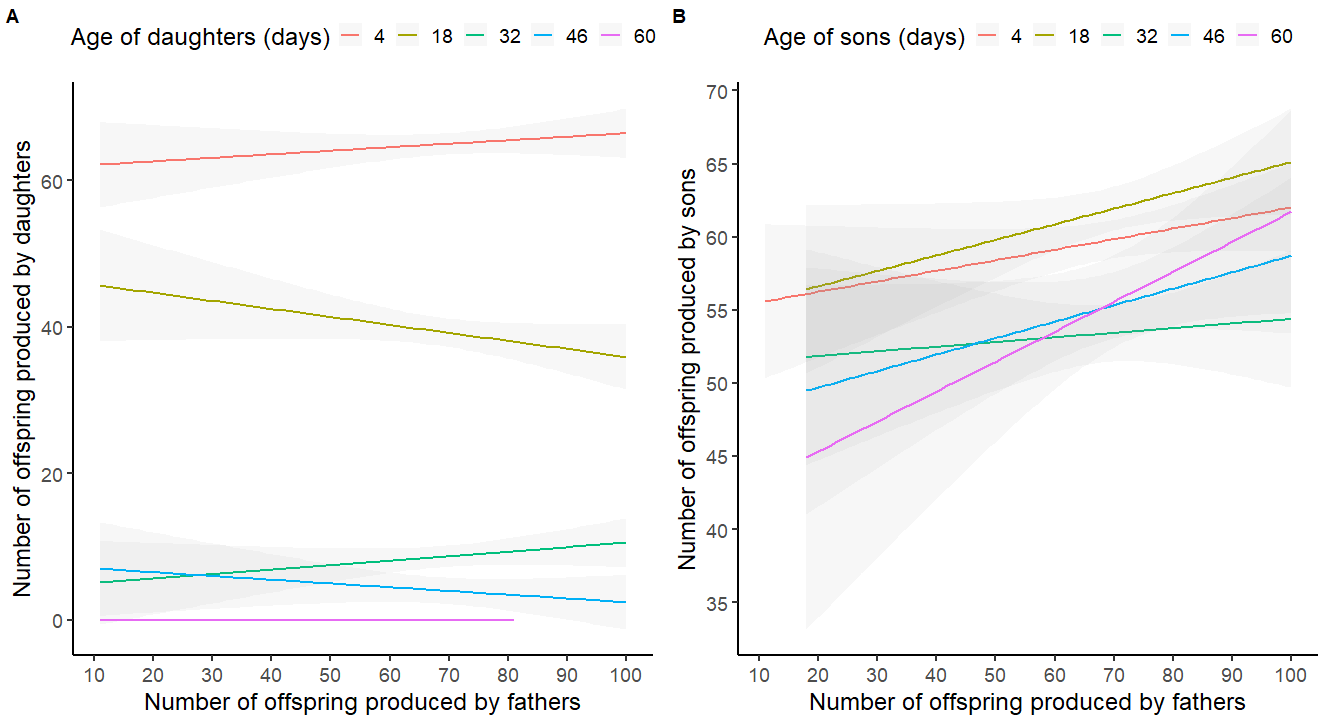

Figure S3. Number of offspring produced by fathers did not affect the number of offspring produced by A) daughters, but had a positive effect on the number of offspring produced by B) sons. Last two age groups (i.e. 74 and 88 days old) were excluded in the figure (but included in the model) for sons, due to very low sample sizes, and to allow better comparison between sons and daughters, given daughters did not live until 74 days.

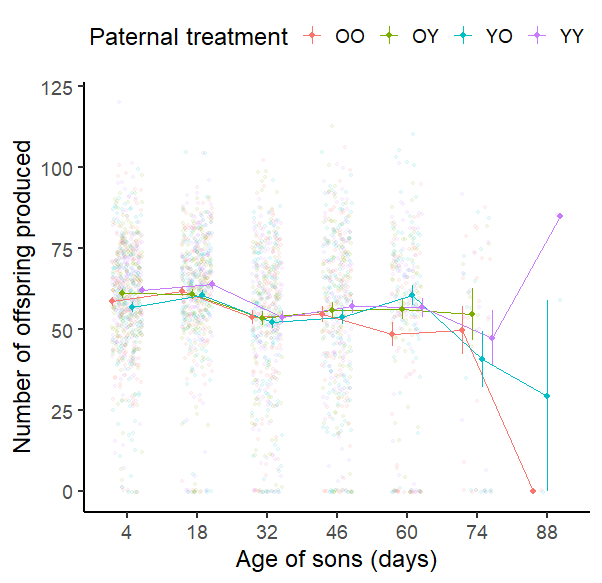

Figure S4: Effect of paternal age and paternal sperm storage duration on the number of offspring produced by sons at different ages. OO: Old paternal age, long paternal sperm storage; OY: Old paternal age, short paternal sperm storage; YO: young paternal age, long paternal sperm storage; YY: young paternal age, short paternal sperm storage. Means and SE along with raw data presented. Each light dot represents measurements on a single son for a given age.

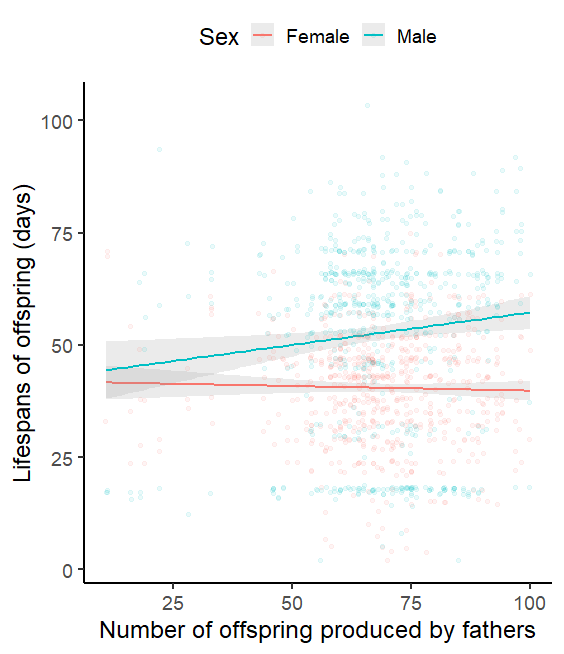

Figure S5: Number of offspring produced by fathers had a significant positive effect on the survival of their sons (male), but not their daughters (female). Lifespans (Y axis), rather than survival probability shown, for simplicity of interpretation, and difficulty of plotting mortality risk hazard function against a continuous variable (paternal reproductive output). Shaded areas show 95% C.I. Each light dot represents measurements on a single offspring.

Appendix 3: Supplementary tables

Table S1: Sample sizes used at each step of our experiment, for each of the four paternal treatments (OO: Old males, long sperm storage; OY: old males, short sperm storage; YO: young males, long sperm storage; YY: young males, short sperm storage). Each row refers to (top to bottom): Number of F0 males measured in the F0 stripping-fertility assay, where each of the 10 females were put on separate vials to assess the proportion of females that produced an offspring; the number of F0 males for whom 10 females from the stripping-counts assay were later kept on individual vials to count the number of offspring produced by each female; the number of fathers mated in the F0 mating assay whose reproductive output was measured; the number of fathers for whom 3 sons and 3 daughters were collected, and later used in the F1 assays.

|  | OO | OY | YO | YY |
| --- | --- | --- | --- | --- |
| N_F0 males stripping-fertility assay | 40 | 34 | 20 | 25 |
| N_F0 males stripping-counts assay | 31 | 22 | 17 | 24 |
| N_F0 fathers mated | 99 | 126 | 101 | 110 |
| N_F1 families | 59 | 60 | 61 | 61 |

Table S2: Effects of age (Y: young, O: old) and sperm storage duration (Y: short, O: long), on number of offspring produced by fathers. Model has zero inflated negative binomial error distribution. Two-way interaction model used to interpret only two-way interaction effect, main effect model used to interpret only main effects. Shaded grey terms were used to interpret associated model effects. Significant terms of interest in bold.

| Two-way interaction model |  |  |  |  |
| --- | --- | --- | --- | --- |
| Dispersion parameter | 7.440 |  |  |  |
| Conditional model: | Estimate | SE | z | P |
| (Intercept) | 4.096 | 0.131 | 31.256 | <0.001 |
| Mating duration | 0.007 | 0.005 | 1.270 | 0.204 |
| RepR2 | -0.112 | 0.056 | -2.004 | 0.045 |
| RepR3 | -0.126 | 0.060 | -2.081 | 0.037 |
| RepR4 | -0.212 | 0.056 | -3.812 | <0.001 |
| Male ageY | 0.081 | 0.062 | 1.297 | 0.195 |
| Sperm storageY | 0.022 | 0.061 | 0.356 | 0.722 |
| Male ageY:Sperm storageY | 0.028 | 0.082 | 0.343 | 0.732 |
| Zero -inflation model | Estimate | SE | z | P |
| (Intercept) | -1.852 | 0.144 | -12.890 | <0.001 |
| Main-effects model |  |  |  |  |
| Conditional model: | Estimate | SE | z | P |
| (Intercept) | 4.088 | 0.129 | 31.670 | <0.001 |
| RepR2 | -0.112 | 0.056 | -2.000 | 0.046 |
| RepR3 | -0.125 | 0.060 | -2.070 | 0.039 |
| RepR4 | -0.212 | 0.056 | -3.810 | <0.001 |
| Mating duration | 0.007 | 0.005 | 1.270 | 0.204 |
| Male ageY | 0.096 | 0.045 | 2.130 | **0.033** |
| Sperm storageY | 0.036 | 0.045 | 0.810 | 0.419 |

Table S3: Effects of paternal age (Y: young, O: old) and paternal sperm storage duration (Y: short, O: long), on age-dependent reproductive output of daughters. Shaded grey terms used to interpret associated model effects. Significant terms of interest in bold. F0_ID = paternal ID, F1_ID = offspring ID.

Three-way interaction model

| Fixed effects | Estimate | SE | z | P |
| --- | --- | --- | --- | --- |
| (Intercept) | 4.182 | 0.087 | 47.860 | <0.001 |
| RepR2 | 0.013 | 0.031 | 0.410 | 0.679 |
| RepR3 | -0.123 | 0.039 | -3.150 | 0.002 |
| RepR4 | -0.004 | 0.040 | -0.100 | 0.920 |
| Daughter’s lifespan | 0.003 | 0.001 | 2.680 | 0.007 |
| Paternal fecundity | 0.000 | 0.001 | 0.040 | 0.969 |
| Paternal ageY | 0.045 | 0.056 | 0.800 | 0.421 |
| Paternal sperm storageY | 0.028 | 0.056 | 0.500 | 0.615 |
| Daughter's age | -0.020 | 0.004 | -5.160 | <0.001 |
| I(Daughter's age^2) | 0.000 | 0.000 | -3.380 | 0.001 |
| Paternal ageY* Paternal sperm storageY | -0.053 | 0.078 | -0.680 | 0.495 |
| Paternal ageY*Daughter's age | -0.004 | 0.003 | -1.260 | 0.209 |
| Paternal sperm storageY*Daughter's age | 0.002 | 0.003 | 0.530 | 0.597 |
| Paternal ageY* Paternal sperm storageY*Daughter's age | 0.001 | 0.005 | 0.300 | 0.761 |
| Random effects | Variance | SD |  |  |
| F0_ID/F1_ID | 0.000 | 0.000 |  |  |
| F0_ID | 0.002 | 0.040 |  |  |

Two-way interaction model

| Fixed effects | Estimate | SE | z | P |
| --- | --- | --- | --- | --- |
| (Intercept) | 4.186 | 0.087 | 48.320 | <0.001 |
| RepR2 | 0.013 | 0.031 | 0.410 | 0.683 |
| RepR3 | -0.123 | 0.039 | -3.150 | 0.002 |
| RepR4 | -0.004 | 0.040 | -0.100 | 0.922 |
| Daughter’s lifespan | 0.003 | 0.001 | 2.690 | 0.007 |
| Paternal fecundity | 0.000 | 0.001 | 0.050 | 0.957 |
| Paternal ageY | 0.035 | 0.046 | 0.770 | 0.442 |
| Paternal sperm storageY | 0.018 | 0.046 | 0.400 | 0.689 |
| Daughter's age | -0.020 | 0.004 | -5.420 | 0.000 |
| I(Daughter's age^2) | 0.000 | 0.000 | -3.450 | 0.001 |
| Paternal ageY*Paternal sperm storageY | -0.035 | 0.049 | -0.700 | 0.481 |
| Paternal ageY*Daughter's age | -0.004 | 0.002 | -1.510 | 0.130 |
| Paternal sperm storageY*Daughter's age | 0.003 | 0.002 | 1.110 | 0.268 |
| Random effects | Variance | SD |  |  |
| F0_ID/F1_ID | 0.000 | 0.001 |  |  |
| F0_ID | 0.002 | 0.040 |  |  |

Main-effects model

| Fixed effects | Estimate | SE | z | P |
| --- | --- | --- | --- | --- |
| (Intercept) | 4.205 | 0.081 | 52.070 | <0.001 |
| RepR2 | 0.012 | 0.031 | 0.400 | 0.689 |
| RepR3 | -0.123 | 0.039 | -3.150 | 0.002 |
| RepR4 | -0.005 | 0.039 | -0.140 | 0.891 |
| Daughter’s lifespan | 0.003 | 0.001 | 2.710 | **0.007** |
| Paternal fecundity | 0.000 | 0.001 | -0.020 | 0.984 |
| Paternal ageY | -0.028 | 0.025 | -1.140 | 0.254 |
| Paternal sperm storageY | 0.035 | 0.024 | 1.430 | 0.153 |
| Daughter's age | -0.021 | 0.003 | -6.050 | **<0.001** |
| I(Daughter's age^2) | 0.000 | 0.000 | -3.410 | **0.001** |
| Random effects | Variance | SD |  |  |
| F0_ID/F1_ID | 0.000 | 0.000 |  |  |
| F0_ID | 0.001 | 0.035 |  |  |

Table S4: Effects of paternal age (Y: young, O: old) and paternal sperm storage duration (Y: short, O: long), on number of offspring produced by sons. Shaded grey terms used to interpret associated model effects. Significant terms of interest in bold. F0_ID = paternal ID, F1_ID = offspring ID.

Three-way interaction model

| Fixed effects | Estimate | SE | z | P |
| --- | --- | --- | --- | --- |
| (Intercept) | 3.850 | 0.085 | 45.570 | <0.001 |
| RepR2 | -0.021 | 0.029 | -0.720 | 0.473 |
| RepR3 | 0.102 | 0.035 | 2.950 | 0.003 |
| RepR4 | 0.123 | 0.035 | 3.570 | <0.001 |
| Son’s lifespan | 0.001 | 0.001 | 1.770 | 0.077 |
| Paternal fecundity | 0.002 | 0.001 | 2.900 | 0.004 |
| Paternal ageY | -0.065 | 0.055 | -1.180 | 0.238 |
| Paternal Sperm storageY | 0.009 | 0.055 | 0.160 | 0.875 |
| Son’s age | -0.004 | 0.001 | -3.420 | 0.001 |
| Paternal ageY* Paternal Sperm storageY | 0.066 | 0.077 | 0.850 | 0.394 |
| Paternal ageY* Son’s age | 0.002 | 0.002 | 1.450 | 0.148 |
| Paternal Sperm storageY * Son’s age | 0.001 | 0.002 | 0.740 | 0.458 |
| Paternal ageY* Paternal Sperm storageY * Son’s age | -0.002 | 0.002 | -0.910 | 0.361 |
| Random effects | Variance | SD |  |  |
| F0_ID/F1_ID | 0.000 | 0.000 |  |  |
| F0_ID | 0.000 | 0.000 |  |  |

Two-way interaction model

| Fixed effects | Estimate | SE | z | P |
| --- | --- | --- | --- | --- |
| (Intercept) | 3.835 | 0.083 | 46.340 | <0.001 |
| RepR2 | -0.021 | 0.029 | -0.720 | 0.469 |
| RepR3 | 0.102 | 0.035 | 2.960 | 0.003 |
| RepR4 | 0.123 | 0.035 | 3.570 | <0.001 |
| Son’s lifespan | 0.001 | 0.001 | 1.740 | 0.083 |
| Paternal fecundity | 0.002 | 0.001 | 2.940 | 0.003 |
| Paternal ageY | -0.036 | 0.045 | -0.800 | 0.422 |
| Paternal Sperm storageY | 0.038 | 0.045 | 0.850 | 0.398 |
| Son’s age | -0.004 | 0.001 | -3.410 | 0.001 |
| Paternal ageY* Paternal Sperm storageY | 0.009 | 0.045 | 0.200 | 0.845 |
| Paternal ageY*Son’s age | 0.001 | 0.001 | 1.130 | 0.257 |
| Paternal Sperm storageY *Son’s age | 0.000 | 0.001 | 0.120 | 0.906 |
| Random effects | Variance | SD |  |  |
| F0_ID/F1_ID | 0.000 | 0.000 |  |  |
| F0_ID | 0.000 | 0.000 |  |  |

Main-effects model

| Fixed effects | Estimate | SE | z | P |
| --- | --- | --- | --- | --- |
| (Intercept) | 3.817 | 0.079 | 48.270 | <0.001 |
| RepR2 | -0.021 | 0.029 | -0.730 | 0.468 |
| RepR3 | 0.101 | 0.035 | 2.940 | 0.003 |
| RepR4 | 0.123 | 0.034 | 3.550 | <0.001 |
| Son’s lifespan | 0.001 | 0.001 | 1.660 | 0.097 |
| Paternal fecundity | 0.002 | 0.001 | 2.930 | **0.003** |
| Paternal ageY | 0.004 | 0.023 | 0.160 | 0.875 |
| Paternal Sperm storageY | 0.046 | 0.023 | 2.030 | **0.043** |
| Son’s age | -0.003 | 0.001 | -4.430 | **<0.001** |
| Random effects | Variance | SD |  |  |
| F0_ID/F1_ID | 0.000 | 0.000 |  |  |
| F0_ID | 0.000 | 0.001 |  |  |

Table S5: Effects of paternal age (Y: young, O: old) and paternal sperm storage duration (Y: short, O: long), on the age-dependent mortality risk for daughters. Shaded terms used to interpret associated model effects. Significant terms of interest in bold. F0_ID = paternal ID.

| Two-way interaction model | coef | exp(coef) | se(coef) | z | P |
| --- | --- | --- | --- | --- | --- |
| RepR2 | -0.503 | 0.605 | 0.120 | -4.190 | 0.000 |
| RepR3 | 0.103 | 1.108 | 0.145 | 0.710 | 0.480 |
| RepR4 | 0.151 | 1.163 | 0.147 | 1.030 | 0.310 |
| Paternal fecundity | 0.002 | 1.002 | 0.003 | 0.500 | 0.610 |
| Paternal ageY | -0.059 | 0.943 | 0.135 | -0.430 | 0.660 |
| Paternal sperm storageY | 0.039 | 1.040 | 0.136 | 0.290 | 0.770 |
| Paternal_ageY*Paternal sperm storageY | -0.004 | 0.996 | 0.190 | -0.020 | 0.980 |
| Random effects | SD | Variance |  |  |  |
| F0_ID | 0.354 | 0.125 |  |  |  |
| Main-effects model | coef | exp(coef) | se(coef) | z | P |
| RepR2 | -0.503 | 0.605 | 0.120 | -4.200 | 0.000 |
| RepR3 | 0.102 | 1.108 | 0.145 | 0.710 | 0.480 |
| RepR4 | 0.151 | 1.162 | 0.147 | 1.030 | 0.310 |
| Paternal fecundity | 0.002 | 1.002 | 0.003 | 0.500 | 0.610 |
| Paternal ageY | -0.061 | 0.941 | 0.096 | -0.630 | 0.530 |
| Paternal sperm storageY | 0.037 | 1.037 | 0.094 | 0.390 | 0.700 |

Table S6: Effects of paternal age (Y: young, O: old) and paternal sperm storage duration (Y: short, O: long), on the age-dependent mortality risk for sons. Shaded grey terms used to interpret associated model effects. Significant terms of interest in bold.

| Two-way interaction model | coef | exp(coef) | se(coef) | z | P |
| --- | --- | --- | --- | --- | --- |
| RepR2 | 0.402 | 1.496 | 0.140 | 2.880 | 0.004 |
| RepR3 | 0.464 | 1.591 | 0.167 | 2.790 | 0.005 |
| RepR4 | 0.168 | 1.183 | 0.170 | 0.990 | 0.320 |
| Paternal fecundity | -0.009 | 0.991 | 0.004 | -2.200 | 0.028 |
| Paternal ageY | 0.066 | 1.068 | 0.156 | 0.430 | 0.670 |
| Paternal sperm storageY | 0.220 | 1.245 | 0.157 | 1.400 | 0.160 |
| Paternal_ageY*Paternal sperm storageY | -0.298 | 0.742 | 0.219 | -1.360 | 0.170 |
| Random effects | SD | Variance |  |  |  |
| F0_ID | 0.539 | 0.290 |  |  |  |
| Main-effects model | coef | exp(coef) | se(coef) | z | P |
| RepR2 | 0.388 | 1.473 | 0.140 | 2.760 | 0.006 |
| RepR3 | 0.454 | 1.574 | 0.168 | 2.700 | 0.007 |
| RepR4 | 0.150 | 1.162 | 0.171 | 0.880 | 0.380 |
| Paternal fecundity | -0.009 | 0.991 | 0.004 | -2.260 | **0.024** |
| Paternal ageY | -0.086 | 0.918 | 0.110 | -0.780 | 0.440 |
| Paternal sperm storageY | 0.065 | 1.067 | 0.110 | 0.590 | 0.550 |
